# Microfluidics on Stretchable Strings

**DOI:** 10.1101/2023.03.02.530606

**Authors:** Philippe DeCorwin-Martin, Andy Ng, David Juncker

## Abstract

This paper introduces string microfluidics (SM), which consists of stretchable strings trapping discrete microdroplets within a porous matrix, and the realization of elementary microfluidic operations such as droplet formation, transport, splitting, merging, and mixing by moving and stretching the strings. While performing these operations, SM was shown to be compatible with colorimetric biological assays. SM represents a new form of microfluidics that integrates the concepts of thread microfluidics and digital microfluidics, along with mobile or reconfigurable microfluidics where liquid transport is realized by manipulating the substrate.

## 1. Introduction

Microfluidics is the manipulation of small volumes of liquids and was initially realized as systems with continuous flow inside microchannels, followed by the development of droplet microfluidics based on the segmentation of flow into discrete droplets. Discrete droplet microfluidics comprises on one hand two-phase flow droplet microfluidics in channels (Thorsen et al., 2001) that allows rapid formation of thousands of droplets and high-throughput biochemical and biological assays, and single-cell analysis (Abatemarco et al., 2017; Guo et al., 2012; Mazutis et al., 2013) but without simple methods for addressing and tracking individual droplets. On the other hand, digital microfluidics uses planar electrodes to electrically actuate droplets which provides addressability (Cho et al., 2003; Miller et al., 2011; Norian et al., 2014), but at the cost of throughput, cost and complexity as advanced electronic are required to actuate and track droplets.

Mobile microfluidics comprises devices with vertical liquid transfer such as pin spotting (Safavieh et al., 2010) or slide-to-slide transfer of droplets arrays such as the ‘SnapChip’(Li et al., 2012; Li et al., 2015). Lateral liquid transfer include the ‘SlipChip’ (Du et al., 2009; Zhukov et al., 2019) that delivers liquid to reconfigurable networks as well as microfluidic probes (Juncker et al., 2005; Qasaimeh et al., 2011; Safavieh et al., 2015). These systems are powerful but require precise alignment and microfabrication.

Capillary microfluidics can operate autonomously and perform complex fluidic operations (Olanrewaju et al., 2018; Olanrewaju et al., 2017; Safavieh and Juncker, 2013; Yafia et al., 2022) without need of any peripheral equipment, and can be made at very low cost using paper (Fu et al., 2011; Giokas et al., 2014; Toley et al., 2015), threads (Rumaner et al., 2019; Safavieh et al., 2011; Tomimuro et al., 2020; Zhou et al., 2012) and flock (Hitzbleck et al., 2013). Threads can be readily functionalized and assembled or knotted into circuits to carry out assays (Weng et al., 2019) and integrated in wearable devices such as skin patches, fabrics and clothing, and wearable devices (Promphet et al., 2020; Xia et al., 2021; Zhao et al., 2021). While fluidic operations in thread-based devices are simple and can be performed through preassembled threads that overlap or tied together, fluid flow on and between the threads are the result of passive wicking, resulting in unreliable fluidic transfer, mixing and retrieval, thus limiting sensitivity and reproducibility.

Herein, we demonstrate a novel mobile and reconfigurable platform for droplet microfluidics. *String microfluidics* (*SM*) consists of stretchable strings that trap droplets within a porous fiber matrix and are used to complete microfluidic operations by active contraction and stretching of the strings to complete fluidic transfer operations. The strings are made of elastic, hydrophilic threads patterned with hydrophobic barriers that isolate the reagent droplets stored in the porous string. Microfluidic operations are performed by pulling, intersecting, twisting, and stretching strings for the transport, transfer, copy, and rapid mixing of droplets. SM represents a new form of microfluidics that integrates the concepts and benefits of (i) digital microfluidics (Choi et al., 2012; Shang et al., 2017; Sohrabi et al., 2020), (ii) thread microfluidics (Agustini et al., 2021; Tan et al., 2021) and (iii) mobile microfluidics where liquid transport is realized by manipulating the substrate (Goyette et al., 2019; Lyu et al., 2019).

## 2. Experimental

### 2.1 String patterning

Strings (Oral-B® Ultra-floss™ and woolly nylon overlock thread) were first washed by sonication in 70% ethanol for 10 min and rinsed in deionized (DI) water. For silanization after washing, the strings were hydrolyzed in 10% hydrochloric acid for 10 min and once again rinsed in DI water. After drying with a nitrogen gun, the strings were silanized with Trichloro(1H,1H,2H,2H-perfluorooctyl)silane (Sigma-Aldrich, Oakville, ON, Canada) for two hours. In the second method, a PAP pen for immunostaining (Sigma-Aldrich, Oakville, ON, Canada) was used that writes with green hydrophobic ink directly creating the barriers once dry. Finally, 4 μL of 1% BSA and 0.05% tween 20 in 1x PBS was spotted onto the strings to create the hydrophilic reservoirs for both patterning methods.

### 2.2 String microfluidics handling

After patterning, the strings were manipulated by hand to achieve transfers, mixing, and copying operations. To facilitate manipulation, the passive strings were taped to holders of various shape or size depending on the application.

### 2.3 Glucose assay

The glucose assay solution consists of a 5:1 solution of glucose oxidase (Sigma-Aldrich, Oakville, ON, Canada) to horseradish peroxidase (Sigma-Aldrich, Oakville, ON, Canada) (150 units of glucose oxidase to 30 units of horseradish peroxidase), 0.6 M potassium iodide, and 0.3 M trehalose at pH 6.0.

## 3. Results and discussion

For the implementation of SM, we selected elastic strings made of curly fibers that straighten when stretched and curl back when loose. In loose strings, the curls in the fibers define large pores, which can accommodate a large volume of liquid. As the string is stretched, the fibers straighten and become closely aligned, reducing the pore size and the volume of the reservoir. These elastic strings constitute an adaptive material as capillary flow can be triggered by stretching the strings (Figure 1a) (Yao et al., 2013). Discrete droplet reservoirs are defined on a stretchable string by forming hydrophobic barriers using fluorosilanes or a hydrophobic marker forming hydrophilic reservoirs that can be loaded with a pipette. Upon stretching the strings, the droplets are drawn out, like liquid being wrung out of a towel (Hadfield, 2013), and remain connected to the hydrophilic fibers by surface tension to form one or several hanging droplets (Figure 1b). When the tension on the string is released, the fibers return to their original, porous conformation, and the liquid is drawn back into the string. Whereas hydrophobic patterning has been used in thread microfluidics to direct fluid flow (Mostafalu et al., 2016), this patterning defines individually addressable reservoirs, enabling easy sample retrieval and manipulation of fluids.

**Figure 1.**
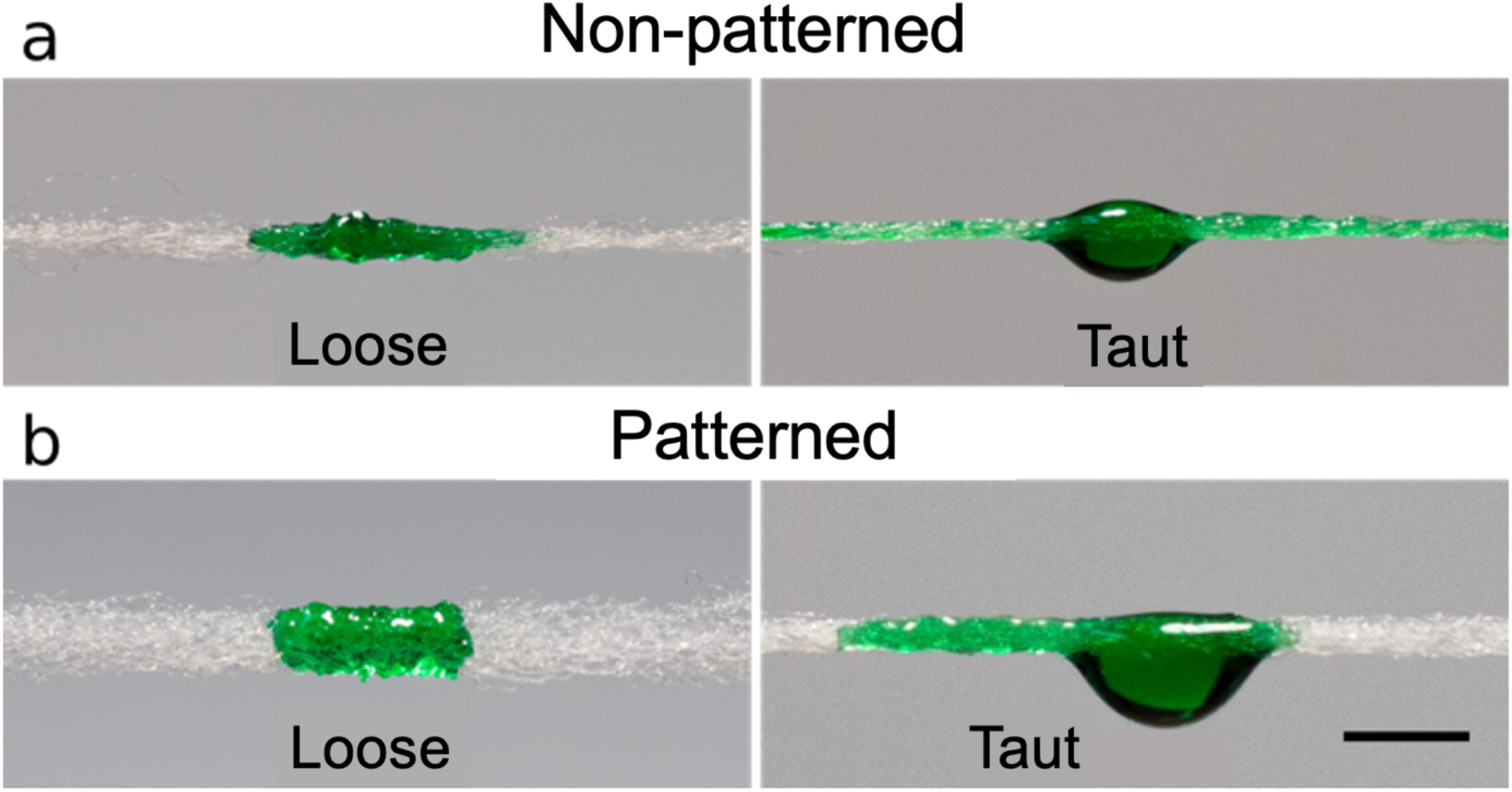
Elastic woolly nylon strings for SM. (a) Non-patterned woolly nylon allows flow along the string when stretched. (b) Patterned woolly nylon extrudes a droplet when stretched, but does not allow flow. Scale bar is 2 mm.

Two strings, Oral-B® Ultra-floss™ and woolly nylon overlock thread were used for SMs. Both strings are made of similar nylon material, but the woolly nylon has a lower fiber count, is thinner, and is more stretchable with a fully stretched strain of ~0.66 (compared to 0.58 for Ultra-floss). Three reservoir states are possible based on the volume of liquid contained: underloaded, over-loaded and operational range. The string is under-loaded when it contains insufficient volume to form beads when the string is stretched. In the case of woolly nylon, this is less than ~0.17 μL mm^-1^ whereas it is slightly higher for Oral-B® Ultra-floss™ as it is 0.5 μL mm^-1^. The operational range for SM manipulations occurs from the minimum volume to form a droplet to volumes of roughly 1 μL mm^-1^ for woolly nylon and 2.1 μL mm^-1^ for Oral-B® Ultrafloss™. Above the operational range, the reservoir becomes over-loaded and a hanging droplet will form in the reservoir, even while the string is loose. When the hanging droplet is large enough, manipulating the string can cause it to fall. It is important to note that the Oral-B^®^ Ultrafloss™ can take larger volumes than the woolly nylon but that both strings have the same functionality and are otherwise used interchangeably. It is expected that one could tune the volumes used in SM by simply adjusting the number of fibers in the string. This would allow smaller or larger volumes to be used depending on the desired application.

Basic SM operations are illustrated in Figure S1 and demonstrated in Figure 2. The transfer of a droplet from a donor to a receiver string can be accomplished by stretching the donor, and contacting it with the loose receiver (Figure 2a, Video S1). The initial volume in the reservoir and the degree to which the string is stretched control the volume transferred. When the donor is fully stretched and contacted to the receiver, the receiver string absorbs all of the droplets and only a minute residue is left on the donor. The mixing of two reagents requires the transfer of an aliquot to a string with a pre-filled reservoir. Since backflow from the receiver to the donor string is a concern, as it would lead to contamination of the donor reservoir, this can be addressed by a gravity-assisted transfer, where the donor string contacts the receiver string from the top for a short time. To demonstrate the limited contamination, a yellow solution was transferred from a donor string to a receiver string filled with a blue solution. The green color on the receiver string highlights successful transfer while the yellow hue retained by the donor string suggests that no significant backflow of blue solution occurred (Figure 2b and Video S2); it is therefore possible to establish unidirectional flow between two droplets on strings.

**Figure 2.**
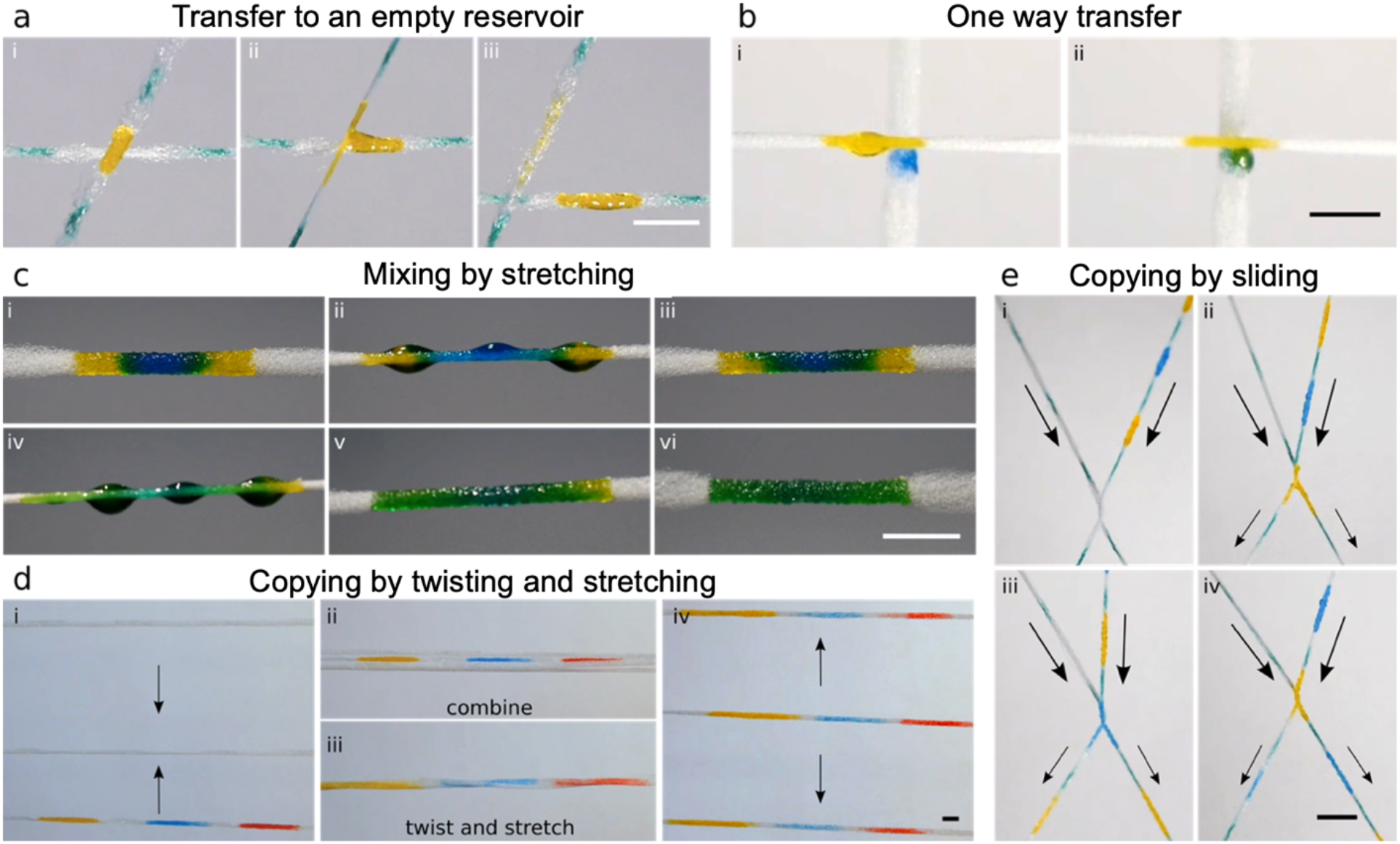
Basic functionality of SMs. (a) Transfer to an empty receiver reservoir. In this transfer, nearly all of the fluid is transferred from the donor to the receiver, leaving only a residue of yellow dye behind, Video S1. (b) One-way reagent transfer of a yellow dye to a receiver string preloaded with a blue dye, Video S2. Contamination of the donor by the blue dye does not occur based on the color integrity. (c) Rapid mixing by cyclic stretching, Video S3. The string is stretched and loosened repeatedly until the reservoir becomes uniform. The procedure was completed in 29 cycles and ~ 25s in this example. (d) Copying by twisting and stretching, Video S5. Initially, a single string has fluid in the reservoirs. To copy the reagents from one string to the others, all strings are combined, twisted, and stretched. Upon separation, all strings are loaded with reagent. (e) Serial copying by sliding, Video S4. The right string’s pattern is transferred to the left string by sliding them past one another. Scale bar is 5 mm.

Following unidirectional transfer, if the droplets do not dry, mixing of the reagents on a string could occur simply by diffusion. The characteristic diffusion time scales as

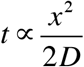

thus, for a reservoir measuring > 1 cm in length and a molecule such as an immunoglobulin G antibody with a diffusion constant of *D_IgG_* = 3.89 ×10^-7^ cm^2^ s^-1^ (Jossang et al., 1988), the time is 357 h, while complete mixing would take even longer. Even for a small molecule, such as fluorescein (*D_f_* = 4.25 × 10^-6^ cm^2^ s^-1^) (Culbertson et al., 2002), the characteristic mixing time is on the order of 33 h. Both mixing times are prohibitively long but are easily reduced using the elasticity of the string to actively mix the reagents by repeatedly stretching the string. Complete mixing of a blue dye transferred to a ~1 cm-long reservoir with a yellow dye was achieved in only 25 s with 29 stretch/release cycles (Figure 2c, Video S3).

A major advantage of SM is that multiple fluidic operations can be achieved by simple manipulations. For example, a common operation in fluidic handling using microwell plates is replicating (or copying), which is done by pipetting reagents from one plate to another. Using SMs, reagents can be rapidly copied onto another string either serially, by sliding two strings across one another (Figure 2e), or in parallel, by twisting multiple receiver strings with a single donor string (Figure 2d). When equal tension is applied to the strings, the fluid is distributed uniformly between the reservoirs. Replicating in series involves contacting the donor and receiver strings and sliding them along their contact point at the same rate such that loaded reservoirs of the donor string come into contact with empty reservoirs of the receiving string one after the other. A transfer rate of one reservoir per second was realized manually. Automation could help increase the transfer rate, but strategies to mitigate string-to-string friction will need to be implemented if considerably higher speeds are desired.

Copying by twisting and stretching allows distributing a liquid to multiple strings simultaneously and is compatible with short strings. Patterned strings, including one with loaded reservoirs, are aligned, brought together, and twisted, then stretched to accomplish copying to all receiving reservoirs simultaneously (Figure 2e, Video S5). We have tested this method using up to 5 woolly nylon receiver strings simultaneously. Above this number, the success rate of copying decreases as there is not enough liquid in the initial reservoir to spread to every string. As it is necessary for the fluid to bead up to bridge the gap and transfer between strings, it is possible to copy one filled reservoir with 1 μL mm^-1^ to a total of 6 equally sized reservoirs for a final volume of 0.17 μL mm^-1^ in each reservoir. However, above this number of stings, each reservoir doesn’t have sufficient liquid to create droplets when stretched and the copying fails. In our experiments, all strings were woolly nylon, but one could easily envision using a thick donor string and transfer it to more than 6 thin receiver strings.

To demonstrate the SM’s compatibility with bio-analysis, a colorimetric glucose assay was performed. This assay is based on a color change produced as iodide is enzymatically oxidized to iodine by horseradish peroxidase (HRP) and glucose oxidase (GOx). For biocompatibility, the support matrix needs to maintain the enzymatic activity of both HRP and GOx as well as ensure that evaporation does not interfere with the reaction. It has previously been used to demonstrate the biochemical compatibility of paper substrate (Martinez et al., 2007), and in development of thread-based biochemical assays (Reches et al., 2010).

A standard curve for the glucose assay was established by patterning one string of three reservoirs with 3 μL of test solution each and then twist copying this donor string to two other receiver strings, resulting in a total of nine test reservoirs. After air drying, a 5 μL aliquot of glucose solution from a dilution series was pipetted to each of the nine glucose test reservoirs and allowed to react for 20 min until dry. The mean RGB intensity was obtained with a desktop scanner and normalized relative to the negative control reservoir (0 mM glucose), yielding a standard curve for the assay (Figure 3). The linear range extends from 0.5 mM to 5 mM, closely matching the results obtained on paper substrates (Martinez et al., 2008). These results indicate that SMs and specifically elastic strings are compatible with enzymatic assays.

**Figure 3.**
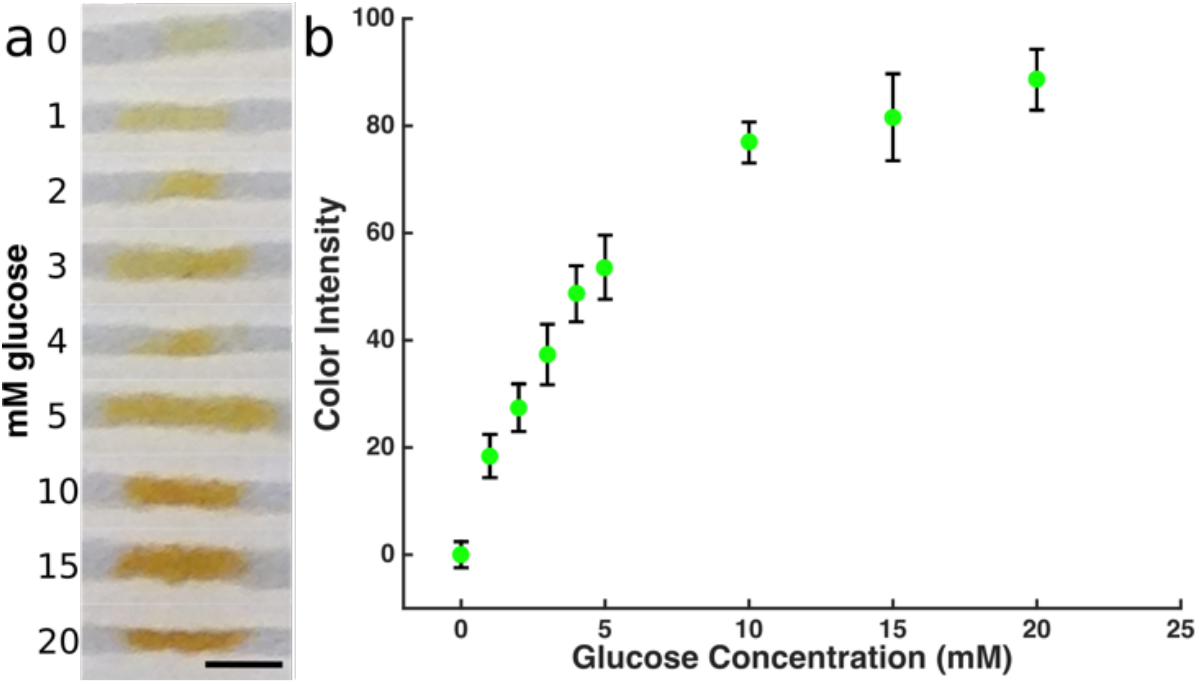
Glucose assay on SMs. (a) Representative color change for glucose dilution series and (b) the color change from three experiments. Error bars represent standard deviation. Scale bar is 3 mm.

## 4. Conclusions

We have introduced SMs which implements the functionality of digital microfluidics on a low-cost platform using porous stretchable threads, hydrophobic barriers to form discrete droplet reservoirs, and physical manipulation by moving and stretching; the stretchable fibrous structure of the string underpins the functionality of SMs. Using SM, arrays of reservoirs can readily be produced and used to conduct biochemical assays. SM stands out by the one-dimensional geometry, and the mechanical actuation scheme.

Flexible strings/threads are building blocks of SM, and form an important class of substrate materials in wearable sensing devices. Strings/threads that are functionalized with enzymes, affinity capture agents and conductive coatings have the capacity of sensing electrolytes, metabolites and biomarkers, and can be integrated in adhesive skin patches, clothing, and other wearable devices (Ates et al., 2022; Weng et al., 2019; Xia et al., 2021). SMs are compatible with reel-to-reel processing for high throughput manufacturing and applications, and can complement other developments towards creating, simple, low cost digital microfluidics, such as paper-based digital microfluidics (Fobel et al., 2014; Ko et al., 2014) or chemically-propelled droplet operations (Cira et al., 2015; Launay et al., 2020). SM enables active fluidic operations such as reagent addition, mixing, and sample retrieval, which can be performed easily by the end-user, and may be combined with other fluidic technologies and materials to establish new microfluidic process flows that take advantage of the unique features of SM. Challenges such as string-to-string cross-contamination, evaporation (and possible re-hydration), will need to be investigated prior to widespread application of SMs. Notwithstanding these challenges, the simplicity of SMs make it an attractive platform to explore linear and random access fluidic operations, and realize complex chemical and biochemical reactions at a low cost.

## Authorship contribution statement

**Philippe DeCorwin-Martin:** Methodology, Data acquisition and curation, Writing — original draft. **Andy Ng:** Writing — original draft, review & editing. **David Juncker:** Funding acquisition, Supervision, Writing —original draft, review & editing.

## Acknowledgement

We acknowledge Grant Ongo for a critical discussion of the manuscript. We acknowledge support from Grand Challenges Canada, NSERC Discovery Grant RGPN-2016-06723. P.D.-M. acknowledges a CGS-M NSERC and a McGill BME Department Recruitment Award. D.J. holds a Canada Research Chair.

## Appendix A. Supplementary data

The following is the Supplementary data to this article:

Supporting figure S1 (PDF)

Video S1: Transfer (MP4)

Video S2: One way transfer (MP4)

Video S3: Mixing (MP4)

Video S4: Copying by Sliding (MP4)

Video S5: Twist mixing (MP4)

## References

Abatemarco, J., Sarhan, M.F., Wagner, J.M., Lin, J.L., Liu, L., Hassouneh, W., Yuan, S.F., Alper, H.S., Abate, A.R., 2017. RNA-aptamers-in-droplets (RAPID) high-throughput screening for secretory phenotypes. Nat Commun 8(1), 332.

Agustini, D., Caetano, F.R., Quero, R.F., Fracassi da Silva, J.A., Bergamini, M.F., Marcolino-Junior, L.H., de Jesus, D.P., 2021. Microfluidic devices based on textile threads for analytical applications: state of the art and prospects. Anal Methods 13(41), 4830–4857.

Ates, H.C., Nguyen, P.Q., Gonzalez-Macia, L., Morales-Narvaez, E., Guder, F., Collins, J.J., Dincer, C., 2022. End-to-end design of wearable sensors. Nat Rev Mater 7(11), 887–907.

Cho, S.K., Moon, H.J., Kim, C.J., 2003. Creating, transporting, cutting, and merging liquid droplets by electrowetting-based actuation for digital microfluidic circuits. J Microelectromech S 12(1), 70–80.

Choi, K., Ng, A.H., Fobel, R., Wheeler, A.R., 2012. Digital microfluidics. Annu Rev Anal Chem (Palo Alto Calif) 5, 413–440.

Cira, N.J., Benusiglio, A., Prakash, M., 2015. Vapour-mediated sensing and motility in two-component droplets. Nature 519(7544), 446–450.

Culbertson, C.T., Jacobson, S.C., Michael Ramsey, J., 2002. Diffusion coefficient measurements in microfluidic devices. Talanta 56(2), 365–373.

Du, W., Li, L., Nichols, K.P., Ismagilov, R.F., 2009. SlipChip. Lab Chip 9(16), 2286–2292.

Fobel, R., Kirby, A.E., Ng, A.H., Farnood, R.R., Wheeler, A.R., 2014. Paper microfluidics goes digital. Adv Mater 26(18), 2838–2843.

Fu, E., Ramsey, S.A., Kauffman, P., Lutz, B., Yager, P., 2011. Transport in two-dimensional paper networks. Microfluid Nanofluidics 10(1), 29–35.

Giokas, D.L., Tsogas, G.Z., Vlessidis, A.G., 2014. Programming fluid transport in paper-based microfluidic devices using razor-crafted open channels. Anal Chem 86(13), 6202–6207.

Goyette, P.A., Boulais, E., Normandeau, F., Laberge, G., Juncker, D., Gervais, T., 2019. Microfluidic multipoles theory and applications. Nat Commun 10(1), 1781.

Guo, M.T., Rotem, A., Heyman, J.A., Weitz, D.A., 2012. Droplet microfluidics for high-throughput biological assays. Lab Chip 12(12), 2146–2155.

Hadfield, C., 2013. Wringing out Water on the ISS - for Science! https://www.youtube.com/watch?v=o8TssbmY-GM.

Hitzbleck, M., Lovchik, R.D., Delamarche, E., 2013. Flock-based microfluidics. Adv Mater 25(19), 2672–2676.

Horning, M.P., Delahunt, C.B., Singh, S.R., Garing, S.H., Nichols, K.P., 2014. A paper microfluidic cartridge for automated staining of malaria parasites with an optically transparent microscopy window. Lab Chip 14(12), 2040–2046.

Jossang, T., Feder, J., Rosenqvist, E., 1988. Photon correlation spectroscopy of human IgG. J Protein Chem 7(2), 165–171.

Juncker, D., Schmid, H., Delamarche, E., 2005. Multipurpose microfluidic probe. Nat Mater 4(8), 622–628.

Ko, H., Lee, J., Kim, Y., Lee, B., Jung, C.H., Choi, J.H., Kwon, O.S., Shin, K., 2014. Active digital microfluidic paper chips with inkjet-printed patterned electrodes. Adv Mater 26(15), 2335–2340.

Launay, G., Sadullah, M.S., McHale, G., Ledesma-Aguilar, R., Kusumaatmaja, H., Wells, G.G., 2020. Self-propelled droplet transport on shaped-liquid surfaces. Sci Rep 10(1), 14987.

Li, H., Bergeron, S., Juncker, D., 2012. Microarray-to-microarray transfer of reagents by snapping of two chips for cross-reactivity-free multiplex immunoassays. Anal Chem 84(11), 4776–4783.

Li, H., Munzar, J.D., Ng, A., Juncker, D., 2015. A versatile snap chip for high-density subnanoliter chip-to-chip reagent transfer. Sci Rep 5, 11688.

Lyu, W., Yu, M., Qu, H., Yu, Z., Du, W., Shen, F., 2019. Slip-driven microfluidic devices for nucleic acid analysis. Biomicrofluidics 13(4), 041502.

Martinez, A.W., Phillips, S.T., Butte, M.J., Whitesides, G.M., 2007. Patterned paper as a platform for inexpensive, low-volume, portable bioassays. Angew Chem Int Ed Engl 46(8), 1318–1320.

Martinez, A.W., Phillips, S.T., Carrilho, E., Thomas, S.W., 3rd, Sindi, H., Whitesides, G.M., 2008. Simple telemedicine for developing regions: camera phones and paper-based microfluidic devices for real-time, off-site diagnosis. Anal Chem 80(10), 3699–3707.

Mazutis, L., Gilbert, J., Ung, W.L., Weitz, D.A., Griffiths, A.D., Heyman, J.A., 2013. Single-cell analysis and sorting using droplet-based microfluidics. Nat Protoc 8(5), 870–891.

Miller, E.M., Ng, A.H., Uddayasankar, U., Wheeler, A.R., 2011. A digital microfluidic approach to heterogeneous immunoassays. Anal Bioanal Chem 399(1), 337–345.

Mostafalu, P., Akbari, M., Alberti, K.A., Xu, Q., Khademhosseini, A., Sonkusale, S.R., 2016. A toolkit of thread-based microfluidics, sensors, and electronics for 3D tissue embedding for medical diagnostics. Microsyst Nanoeng 2(1), 16039.

Norian, H., Field, R.M., Kymissis, I., Shepard, K.L., 2014. An integrated CMOS quantitative-polymerase-chain-reaction lab-on-chip for point-of-care diagnostics. Lab Chip 14(20), 4076–4084.

Olanrewaju, A., Beaugrand, M., Yafia, M., Juncker, D., 2018. Capillary microfluidics in microchannels: from microfluidic networks to capillaric circuits. Lab Chip 18(16), 2323–2347.

Olanrewaju, A.O., Ng, A., DeCorwin-Martin, P., Robillard, A., Juncker, D., 2017. Microfluidic Capillaric Circuit for Rapid and Facile Bacteria Detection. Anal Chem 89(12), 6846–6853.

Promphet, N., Hinestroza, J.P., Rattanawaleedirojn, P., Soatthiyanon, N., Siralertmukul, K., Potiyaraj, P., Rodthongkum, N., 2020. Cotton thread-based wearable sensor for non-invasive simultaneous diagnosis of diabetes and kidney failure. Sensors and Actuators B: Chemical 321.

Qasaimeh, M.A., Gervais, T., Juncker, D., 2011. Microfluidic quadrupole and floating concentration gradient. Nat Commun 2, 464.

Reches, M., Mirica, K.A., Dasgupta, R., Dickey, M.D., Butte, M.J., Whitesides, G.M., 2010. Thread as a matrix for biomedical assays. ACS Appl Mater Interfaces 2(6), 1722–1728.

Rumaner, M., Horowitz, L., Ovadya, A., Folch, A., 2019. Thread as a Low-Cost Material for Microfluidic Assays on Intact Tumor Slices. Micromachines (Basel) 10(7).

Safavieh, M., Qasaimeh, M.A., Vakil, A., Juncker, D., Gervais, T., 2015. Two-Aperture Microfluidic Probes as Flow Dipole: Theory and Applications. Sci Rep 5, 11943.

Safavieh, R., Juncker, D., 2013. Capillarics: pre-programmed, self-powered microfluidic circuits built from capillary elements. Lab Chip 13(21), 4180–4189.

Safavieh, R., Roca, M.P., Qasaimeh, M.A., Mirzaei, M., Juncker, D., 2010. Straight SU-8 pins. J Micromech Microeng 20(5), 055001.

Safavieh, R., Zhou, G.Z., Juncker, D., 2011. Microfluidics made of yarns and knots: from fundamental properties to simple networks and operations. Lab Chip 11(15), 2618–2624.

Shang, L., Cheng, Y., Zhao, Y., 2017. Emerging Droplet Microfluidics. Chem Rev 117(12), 7964–8040.

Sohrabi, S., Kassir, N., Keshavarz Moraveji, M., 2020. Droplet microfluidics: fundamentals and its advanced applications. RSC Adv 10(46), 27560–27574.

Tan, W., Powles, E., Zhang, L., Shen, W., 2021. Go with the capillary flow. Simple thread-based microfluidics. Sensors and Actuators B: Chemical 334.

Thorsen, T., Roberts, R.W., Arnold, F.H., Quake, S.R., 2001. Dynamic pattern formation in a vesicle-generating microfluidic device. Phys Rev Lett 86(18), 4163–4166.

Toley, B.J., Wang, J.A., Gupta, M., Buser, J.R., Lafleur, L.K., Lutz, B.R., Fu, E., Yager, P., 2015. A versatile valving toolkit for automating fluidic operations in paper microfluidic devices. Lab Chip 15(6), 1432–1444.

Tomimuro, K., Tenda, K., Ni, Y., Hiruta, Y., Merkx, M., Citterio, D., 2020. Thread-Based Bioluminescent Sensor for Detecting Multiple Antibodies in a Single Drop of Whole Blood. ACS Sens 5(6), 1786–1794.

Weng, X., Kang, Y., Guo, Q., Peng, B., Jiang, H., 2019. Recent advances in thread-based microfluidics for diagnostic applications. Biosens Bioelectron 132, 171–185.

Xia, J., Khaliliazar, S., Hamedi, M.M., Sonkusale, S., 2021. Thread-based wearable devices. MRS Bulletin 46(6), 502–511.

Yafia, M., Ymbern, O., Olanrewaju, A.O., Parandakh, A., Sohrabi Kashani, A., Renault, J., Jin, Z., Kim, G., Ng, A., Juncker, D., 2022. Microfluidic chain reaction of structurally programmed capillary flow events. Nature 605(7910), 464–469.

Yao, X., Hu, Y., Grinthal, A., Wong, T.S., Mahadevan, L., Aizenberg, J., 2013. Adaptive fluid-infused porous films with tunable transparency and wettability. Nat Mater 12(6), 529–534.

Zhao, C., Li, X., Wu, Q., Liu, X., 2021. A thread-based wearable sweat nanobiosensor. Biosens Bioelectron 188, 113270.

Zhou, G., Mao, X., Juncker, D., 2012. Immunochromatographic assay on thread. Anal Chem 84(18), 7736–7743.

Zhukov, D.V., Khorosheva, E.M., Khazaei, T., Du, W., Selck, D.A., Shishkin, A.A., Ismagilov, R.F., 2019. Microfluidic SlipChip device for multistep multiplexed biochemistry on a nanoliter scale. Lab Chip 19(19), 3200–3211.

