## Supplementary information for "Microfluidics on Stretchable Strings"

### Supporting Information

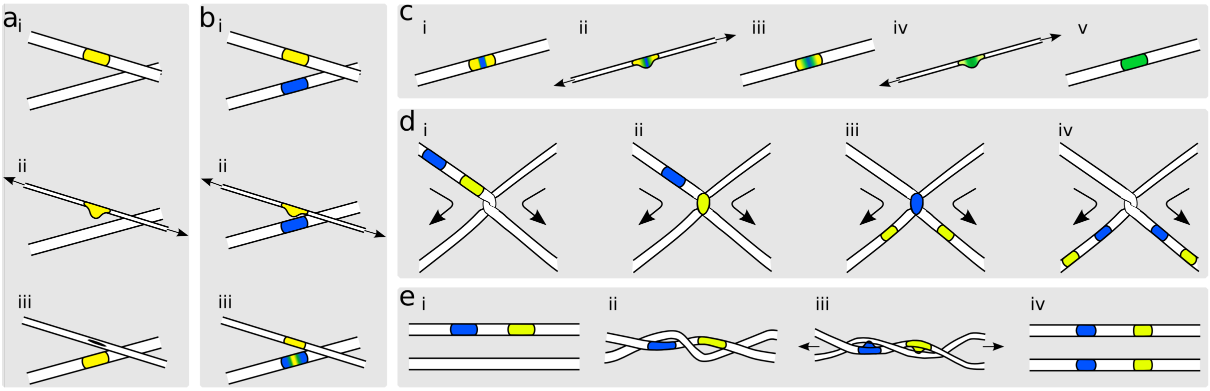


**Figure S1.** Schematic of the five basic principles and fluidic functions realized by SMs. One-way transfer of reagents from donor to receiver string with (a) an empty and (b) a pre-loaded receiver string. (c) Active, rapid mixing of two reagents by cyclic stretching. Rapid copying of reagents from a string to one or multiple other strings by (d) sliding and (e) twisting and stretching.
